# Human iPS Cells Derived Skeletal Muscle Progenitor Cells Promote Myoangiogenesis and Restore Dystrophin in Duchenne Muscular Dystrophic Mice

**DOI:** 10.1101/2020.01.21.914283

**Authors:** Wanling Xuan, Mahmood Khan, Muhammad Ashraf

**Affiliations:** Vascular Biology Center, Medical College of Georgia at Augusta University, Augusta, Georgia, USA; Department of Medicine, Medical College of Georgia at Augusta University, Augusta, Georgia, USA; Department of Emergency Medicine, Wexner Medical Center, The Ohio State University, Columbus, Ohio, USA

**Keywords:** Duchenne Muscular Dystrophy, Human induced pluripotent stem cells, muscle progenitor cells, histone deacetylase inhibitor, angiogenesis

## Abstract

**Background and Objective:** Duchenne muscular dystrophy (DMD) is caused by mutations of the gene that encodes the protein dystrophin. Loss of dystrophin leads to severe and progressive muscle-wasting in both skeletal and heart muscles. Human induced pluripotent stem cells (hiPSCs) and their derivatives offer important opportunities to treat a number of diseases. Here, we investigated whether givinostat, a histone deacetylase inhibitor (HDACi), could reprogram hiPSCs into muscle progenitor cells (MPC) for DMD treatment.

**Methods and Results:** MPC generated by CHIR99021 and givinostat (Givi) small molecules from multiple hiPSCs expressed myogenic makers (Pax7, desmin) and were differentiated into myotubes expressing MF20 upon culture in specific differentiation medium. These MPC exhibited superior proliferation and migration capacity determined by CCK-8, colony and migration assays compared to control-MPC generated by CHIR99021 and fibroblast growth factor (FGF). Upon transplantation in hind limb of Mdx/SCID mice with cardiotoxin (CTX) induced injury, these MPC showed higher engraftment and restoration of dystrophin than treatment with control-MPC and human myoblasts. In addition, treated muscle with these MPC showed significantly limited infiltration of inflammatory cells and reduced muscle necrosis and fibrosis. A number of these cells were engrafted under basal lamina expressing Pax7, which were capable of generating new muscle fibers after additional injury. Extracellular vesicles released from these cells promoted angiogenesis after reinjury.

**Conclusion:** We successfully generated integration free MPC from multiple hiPS cell lines using CHIR99021 and Givi. Givinostat induced MPC showed marked and impressive regenerative capabilities and restored dystrophin in injured tibialis muscle compared to control MPC. Additionally, MPC generated by Givi also seeded the stem cell pool in the treated muscle. It is concluded that hiPSCs pharmacologically reprogrammed into MPC with a small molecule, Givi with anti-oxidative, anti-inflammatory and muscle gene promoting properties might be an effective cellular source for treatment of muscle injury and restoration of dystrophin in DMD.

## Introduction

Duchenne muscular dystrophy (DMD) is caused by mutations of the gene that encodes the protein dystrophin. Loss of dystrophin leads to severe and progressive muscle-wasting in both skeletal and heart muscles. Cell replacement gives a promising hope for DMD therapy. Satellite cells (SCs) are endogenous skeletal muscle stem cells, which are responsible for muscle maintenance and muscle regeneration after injury (1, 2). A previous study reported that xenotransplantation of human SCs into mice achieved efficient engraftment and populated the satellite niche (3). However, a biopsy is needed for procurement of SCs. In addition, freshly isolated SCs progeny though can be propagated *in vitro* but their transplantation potential becomes limited during *in vitro* expansion (4–6). Therefore, procurement of larger number of SCs for transplantation becomes an obstacle for clinical application. Human induced pluripotent stem cells (hiPSCs) derived derivatives offer important sources to treat a number of diseases. Efforts have been made in the past few years for generation of muscle progenitor cells (MPC) from hiPSCs either by genetic modification or small molecules. Nevertheless, generation of MPC from hiPSCs by viral vectors remains a safety concern. High percentage of Pax7 positive MPC can be generated from hiPSC by small molecules (CHIR99021, LDN19389 and FGF) (7, 8), but their limited engraftment was observed *in vivo* upon transplantation (9). Interestingly, it has been recently reported that MPC can be generated from teratoma which showed high engraftment efficiency in muscle dystrophy model (10). However, human teratoma derived MPC poses safety concerns for clinical application. Therefore, it seems more appropriate to look for alternate approaches for inducing MPC from hiPSCs with high engraftment and differentiation properties.

Givinostat is a histone deacetylase inhibitor (HDACi) that has been shown to increase muscle regeneration in a mouse model of DMD (11). Interestingly, most of the beneficial effects of HDACi arise from its ability to redirect fibroadipogenic lineage commitment toward a myogenic fate (12). Using genome-wide Chip-seq analysis in myoblasts, it was demonstrated that HDACi induced myogenic differentiation program in myoblasts (i.e., Myosin 7, Enolase 3 and Myomesin1) (13). Therefore, here we propose that Givi could reprogram hiPSCs into MPC for DMD treatment.

## Methods

### Human iPSC culture

The Human iPSC cell lines from ATCC Company CYS0105 and DYS0100 were used. CYS0105 was reprogrammed from human cardiac fibroblasts of a 72 years old healthy donor, while DYS0100 was reprogrammed from human foreskin fibroblasts of a normal newborn. DMD-iPS cell line (SC604A MD) was purchased form SBI Company, which was generated from a DMD patient with Exon 3-7 deletion of dystrophin. The forth iPS cell line was reprogrammed from human dermal fibroblasts (CC-2511, Lonza) of a 45 years old healthy donor in our lab using Cyto TuneTM iPS 2.0 sendai reprogramming kit (A16517, Thermo fisher Scientific) as previously described (14). iPSCs were grown and maintained on vitronectin coated six-well plate in mTeSR1 medium (Stem Cell Technologies) with daily change.

### Differentiation protocols to generate muscle progenitor cells (MPC) and their characterization

Human iPSCs at passage 20-30 were used for conversion to MPC. Human iPSCs were dissociated into single cells using Accutase (Stem Cell Technologies) at 37℃ for 10 min and then were seeded on to a vitronectin-coated six-well plate at 3×10^5^ cell/well in mTeSR1 supplemented with 5μM ROCK inhibitor (Y-27632, Stem Cell Technology) for 24 h. Afterwards cells were switched into E6 medium (Thermo Fisher Scientific) supplemented with CHIR99021 (10 μM) for two days followed by Givi (100 nM) for 5 days. The differentiating cells were cultured in E6 medium for 7 days. The schematic outline is shown in Fig.1A. At Day 14, cells were replated on 0.1% galectin coated coverslips and expression of Pax7 and desmin were analyzed by immunostaining. MPC were expanded in SKGM-2 medium plus FGF-2 (2.5 ng/ml) and cells at passage 2-4 were used for experiments. Here, we referred the givinostat induced MPC as Givi-MPC. To further enhance muscle differentiation, at Day 14, cultured cells were replated and switched into high glucose DMEM medium supplemented with 2% horse serum (Thermo fisher Scientific) and 1% ITS (Thermo fisher Scientific) for 7 days. Immunostaining of cultured cells for MF20 was performed. To generate control-MPC (7), a previously reported method using only CHIR99021 was used. The schematic outline is shown in supplemental **Fig.S1**. Control-MPC were expanded using the same condition as Givi-MPC and passage 2-4 of cells were used for experiments.

**Fig. 1.**
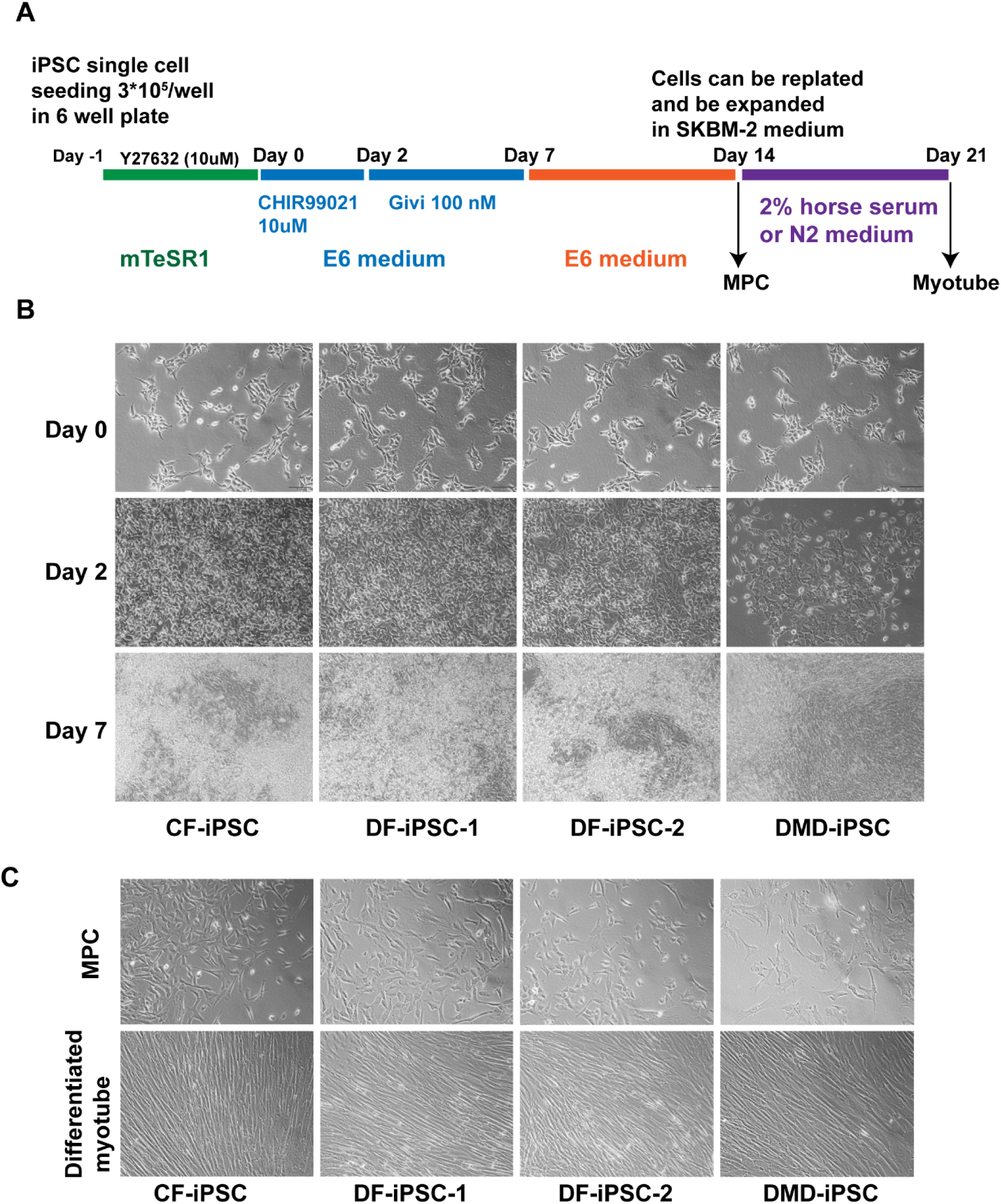
Generation of muscle progenitor cell (MPC) from human iPSC using small molecules. (A) Schematic outline of generation of MPC from human PSC using combination of CHIR99021 and givinostat or CHIR99021 only. (B) Morphology of differentiating cells from 4 human iPSC lines (CF-iPSC, DF-iPSC-1, DF-iPSC-2 and DMD-iPSC) at 7 days. Bar = 200 μm. (C) Morphology of replated MPC and differentiated myotubes from 4 human iPSC lines at day 14. Bar = 200 μm.

### CCK-8 assay for proliferation

CCK-8 assay was used for evaluation of cell proliferation. Briefly, 2000 cells were seeded into 96 well plate per well and cell proliferation was analyzed at 0h, 24h, 48h and 72h respectively by using CCK-8 kit (ab228554, abcam) according to the manufacturer protocol.

### Colony formation

Thirty cells (single cell) were seeded in one well of six-well plate. After 7 days, cells were stained with crystal violet dye. Number of colonies and size of cell growth were analyzed and compared between control-MPC and Givi-MPC groups.

### Cell migration

For cell migration experiment, human myoblasts, control-MPC and Givi-MPC were seeded in 35mm dish with culture-insert 2 well (ibidi company) at 1×10^5^/ml concentration in SKGM-2 medium with 2% Fetal Bovine Serum (FBS). The next day, a confluent layer was observed and culture-inserts were removed, and after 24h the number of migrated cells were analyzed.

### Human endothelial cell and human myoblast culture

Human aortic endothelial cells (HAEC, CC-2535) and human skeletal myoblasts (HSMM-Muscle Myoblasts, CC-2580) were obtained from Lonza Company. HAEC were maintained in endothelial cell growth medium V-2 (213-500, CELL APPCLICATIONS, Inc.) and cells at passage 2-6 were used for experiments. Human myoblasts were maintained in SKGM-2 medium (Lonza) and cells at passage 2-4 were used for experiments.

### Cardiotoxin injury and cell transplantation

Animal experiments were carried out according to experimental protocol approved by the Augusta University Animal Care and Use Committee. 6-8 weeks old Mdx/SCID mice (Stock No: 018018, The Jackson Laboratory) were used in the present study. One-day prior to cell transplantation, mice were anaesthetized using 2% isoflurane and tibialis anterior (TA) muscle was injured with 50 μl of 10 μM cardiotoxin (Naja mossambica-mossambica, Sigma). For cell transplantation experiments, control-MPC and Givi-MPC were differentiated from the same human iPS cell line, DYS0100. For transplantation, myoblast, control-MPC and Givi-MPC were dissociated using Accutase (Stem Cell Technologies) and resuspended in Dulbecco’s phosphate-buffered saline (DPBS) at 1×10^5^ per 20 μl. Cells were injected into the left TA muscle while the same volume of DPBS was injected into the right TA as control. In some cases, cells were transfected with Green Fluorescent Protein (GFP) Lentivirus (abm company, Canada) for cell tracking. Some Mdx/SCID mice transplanted with Givi-MPC were subjected to CTX reinjury at 2M after first injury and cell transplantation.

### Immunostaining for cells

Cells were fixed with 4% PFA, and blocked with 10% FBS, followed by incubation with anti-Pax7 antibody (ab187339, abcam, 1:300), anti-desmin antibody (ab32362, abcam, 1:500) and anti-Myosin Heavy Chain Antibody (MF20) antibody (Novus, MAB4470, 1:200) respectively at 4°C overnight and secondary antibody conjugated to Alexa Fluor 594 or Alexa Fluor 488 (Life Technologies) at room temperature for 1h. Images were taken by a florescent microscope (Olympus, Japan).

### Immunostaining for muscle sections

After 7 days or 30 days of cell transplantation, Mdx/SCID mice were euthanized and TA muscles were harvested and fixed with 4% paraformaldehyde (PFA) for 1h at room temperature and then immersed in 30% sucrose overnight at 4°C. At day two, hearts were cryopreserved in an optical cutting temperature (OCT) compound (Tissue Tek) at −80°C. TA muscle samples were sliced into 5-μm-thick frozen cross-sections using a Leica CM3050 cryostat. Muscle sections were incubated with primary antibodies including Laminin (L9393, Sigma,1:500), dystrophin (D8168, Sigma, 1:200), human specific laminin (LAM-89, Novus, 1:200), GFP (#2956, Cell Signal Technologies, 1:500), dystrophin (ab15277, abcam, 1:200), human nuclear antigen (NBP2-34342, Novus, 1:100), CD68 (NB600-985, Novus, 1:200) and CD31 (NB600-562, Novus, 1:200) at 4°C overnight respectively and anti-rabbit/mouse secondary antibodies conjugated to Alexa Fluor 594 or Alexa Fluor 647 or Alexa Fluor 488 (Life Technologies) at room temperature for 1h. Images were taken using a confocal microscope (FV1000, Olympus, Japan). For cell engraftment quantification, 4 sections at 150 µm interval in each TA muscle were analyzed. Dystrophin or laminin staining was used to define the physical boundaries of muscle fibers. The number of muscle fibers and cross-section area were measured using Image J with the colocalization plugin (NIH). Capillary density was assessed in 4 sections cut at 150 µm interval by counting CD31 positive vascular structures using a fluorescence microscope at a magnification of 400 x. The number of capillaries in each TA muscle was expressed as the number of capillary per field (0.2 mm^2^). For quantification of inflammatory cells, number of CD68 positive cells were counted in 3 sections cut at 150 µm interval after 7 days’ post cell transplantation and was expressed as the number of CD68 positive cells per field (0.2 mm^2^). Staining of presynaptic marker α-bungarotoxin (α-BTX) was carried out using α-bungarotoxin, Alexa Fluor™ 594 conjugate (Invitrogen) according to the manufacturer’s instruction.

### Histology

Histological staining was performed at Electron Microscopy and Histology Core of Augusta University. After 7 days or 30 days of cell transplantation, TA muscle were harvested and embedded in paraffin. 5-μm-thick sections of TA muscle were cut and stained with hematoxylin and eosin (H and E), Masson trichrome and Sirius red according to the manufacturer protocol (abcam). Images were taken by a vertical microscope (Olympus, Japan). Fibrosis and necrosis were determined using the ImageJ software (NIH) and expressed as the ratio of total area of the cross-section and normalized with the ratio of control lateral TA muscle section. Myofiber necrosis was identified with fragmented sarcoplasm (15) and/or increased inflammatory cell infiltration, and was measured using non-overlapping tile images of transverse muscle sections that provided a view of the entire muscle cross section.

### Isolation of extracellular vesicles (EV)

EV were isolated using size exclusion column method as we described previously (16). Briefly, conditioned media was collected and EV were isolated by centrifugation at 3000 rpm for 30 min to remove cells and debris, followed by filtration through a 0.22 μm filter to remove the remaining debris. Then the medium was further concentrated using Amicon Ultra-15 100 KDa centrifugal filter units (Millipore). Isolation of EV in the concentrated medium was carried out through qEV size exclusion columns (Izon Science). EV fractions were collected and concentrated by Amicon Ultra-4 10 KDa centrifugal filter (Millipore). The purified EV were stored at −80°C and subsequently characterized by particle size and electron microscopy.

### Concentration and Particle size measurement with tunable resistive pulse sensing

Particle size and concentration were analyzed using tunable resistive pulse sensing (TRPS) technique with a qNano instrument (Izon Science) as described in previous studies (16, 17). Briefly, the number of particles were counted (at least 600 to 1000 events) at 20 mbar pressure. Beads CPC200 (200 nm) were used for calibration. Data were analyzed using Izon Control Suite software.

### Transmission electron microscopy (TEM)

Tissue samples were processed for TEM by the Electron Microscopy and Histology Core Laboratory at Augusta University as described previously (16). Briefly, EV suspension was fixed with an equal volume of 8% paraformaldehyde to preserve ultrastructure. Ten µl of suspended/fixed exosomes was applied to a carbon-formvar coated 200 mesh copper grid and allowed to stand for 30-60 seconds. The excess was absorbed by Whatman filter paper. 10 µl of 2% aqueous uranyl acetate was added and treated for 30 seconds. Grids were allowed to air dry before being examined in a JEM 1230 transmission electron microscope (JEOL USA Inc., Peabody, MA) at 110 kV and imaged with an UltraScan 4000 CCD camera & First Light Digital Camera Controller (Gatan Inc., Pleasanton, CA).

### RNA extraction and PCR array

Total RNA from cells was isolated using miRNeasy Kit (Qiagen). Reverse transcription was performed using QuantiTect Reverse Transcription kit (Qiagen). Human cell motility RT2 profiler PCR Array (Qiagen) for control-MPC and Givi-MPC was performed. Data was analysed using RT2 Profiler PCR Array Data Analysis Webportal (Qiagen). Genes with a fold change >2.0 were considered overexpressed.

### RNA extraction from EV and miRNA Array analysis

Total RNA from EV was isolated using miRNeasy Micro Kit (Qiagen). The miRNA Array analysis was performed in the Integrated Genomics and High Performance Computer Server center at Augusta University. RNA purity and concentration were evaluated by spectrophotometry using Nanodrop ND-1000 (Thermo Fisher Scientific). Quality and the related size of small RNA was assessed by the Agilent 2100 Bioanalyzer (Agilent Technologies, Santa Clara, CA). 130 ng of total RNA was labeled with biotin using the FlashTag Biotin HSR RNA Labeling Kit (Applied Biosystems) according to manufacturer’s procedure. The labeled samples were then hybridized to the GeneChip miRNA 4.0 array (Thermofisher) that contains 2,578 and 2,025 human mature and premature miRNA, respectively. Array hybridization, washing, and scanning of the arrays were carried out according to Affymetrix’s recommendations. Data was obtained in the form of CEL file. The CEL files were imported into Partek Genomic Suites version 6.6 (Partek, St. Louis, MO) using standard import tool with RMA normalization. The differential expressions were calculated using ANOVA of Partek Package.

### Tube formation assay

Human aortic endothelia cells (HAEC, 1×10^5^ cells/well) were seeded on Matrigel (Corning) in a 24-well plate and treated with or without 1 μg EV from different groups of Givi-MPC, control-MPC and human myoblast in EGM-2V basal medium (Lonza). After 16 h, cells in Matrigel were stained with Calcein AM, and images were taken with fluorescent microscope. Tube formation was analysed by Image J software with the angiogenesis analyzer plugin (NIH).

### Statistical analysis

Data were expressed as mean ± SD. After test for normality, statistical analysis of differences among different groups was compared by ANOVA with Bonferroni’s correction for multiple comparisons. Percentage of different size of colony was compared using Chi-squared test. The Differences were considered statistically significant at *P < 0.05*. Statistical analyses were performed using Graphpad Prism 6.0 (Chicago, US).

## Results

### Generation of muscle progenitor cells from human iPSC using small molecules

As outlined in Fig.1A, we used 3 iPSC lines from healthy donors with different ages, and one iPSC line from DMD patient with frameshift deletions of exons 3-7 in the dystrophin gene for MPC generation. After 2 days treatment with CHIR99021, the morphology of the differentiating cells from 4 cell lines was dramatically altered indicative of epithelial to mesenchymal transition (EMT) (**Fig.1B**). Following treatment with givinostat for 5 days, cells became confluent and clustered (**Fig.1B**). Fig.1C showed the morphology of differentiated MPC after replating and terminal muscle differentiation for 7 days in 2% horse serum differentiation medium. Using immunostaining, the MPC derived from 4 iPSC lines expressed the myogenic markers Pax7 and desmin. The MPC during terminal differentiation exhibited elongated shape (**Fig.1C**) and expressed MF20 (**Fig.2B**), indicating their myogenic differentiation potential.

**Fig. 2.**
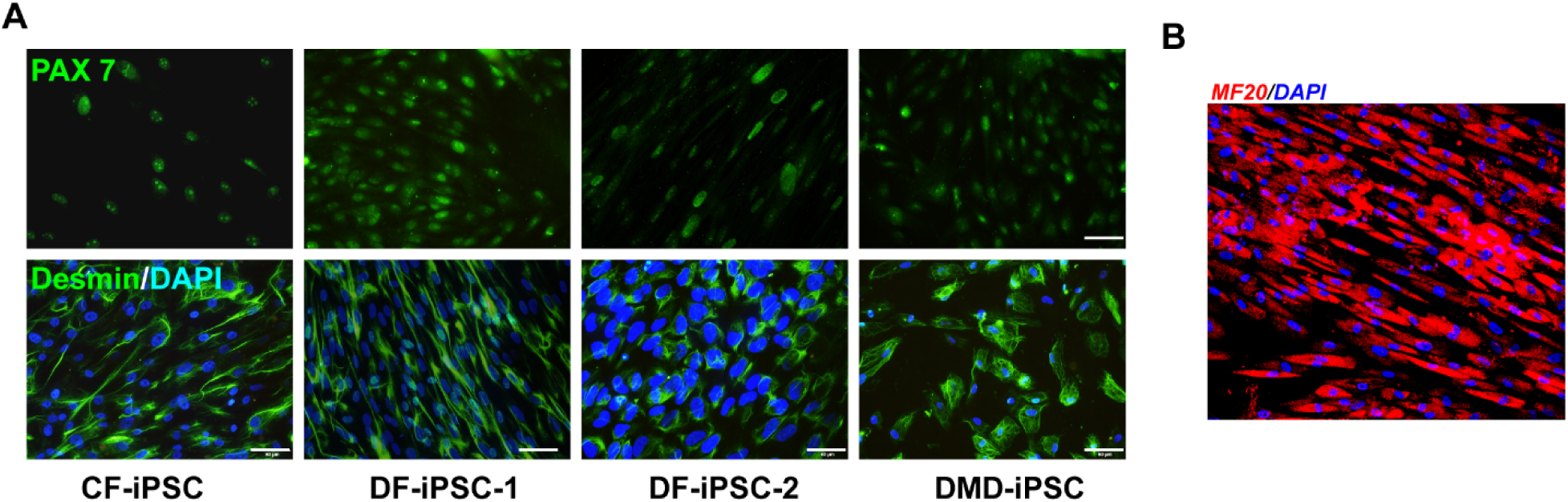
Characterization of givinostat-induced MPC. (A) The treated hiPSC at day14 expressed Pax7 and desmin. (B) The differentiated myotubes expressed MF20 as shown by immunostaining. Bar = 50 μm.

### Givinostat induced MPC (Givi-MPC) expressed high proliferation and motility properties *in vitro*

Next, we explored whether MPCs were proliferative and possessed self-renewal and motility properties. Using migration assay, compared to normal adult human myoblasts or control-MPC, Givi-MPC exhibited superior migration capability in low serum medium culture (**Fig.3A**) with highest number migrated compared to other MPCs (**Fig.3B**). Genes related to migration as determined by cell motility PCR array increased multifold in Givi-MPC as compared with MPCs (ITGA4 (25.61 fold), RAC2 (7.48 fold), FGF2 (6.75 fold) AND ENAH (5.53 fold)). **Fig.3C** and **Fig.3D** show heatmap and the list of upregulated genes related to migration (> 2 fold). In addition, CCK-8 assay at 48h and 72h time points Givi-MPC showed higher OD value compared with human myoblasts and control-MPC (**Fig.3E**). These data suggest that Givi-MPC possess better self-renewal potential. Colonies formed by Givi-MPC were bigger and had a higher cell density compared to control-MPC (**Fig.3F**). Quantitative data (**Fig.3G** **and 3H**) showed that Givi-MPC formed more colonies with higher density of cells (>200 cells) compared to control-MPC. These data support the notion that Givi-MPC possessed highly proliferation and self-renewal capabilities.

**Fig. 3.**
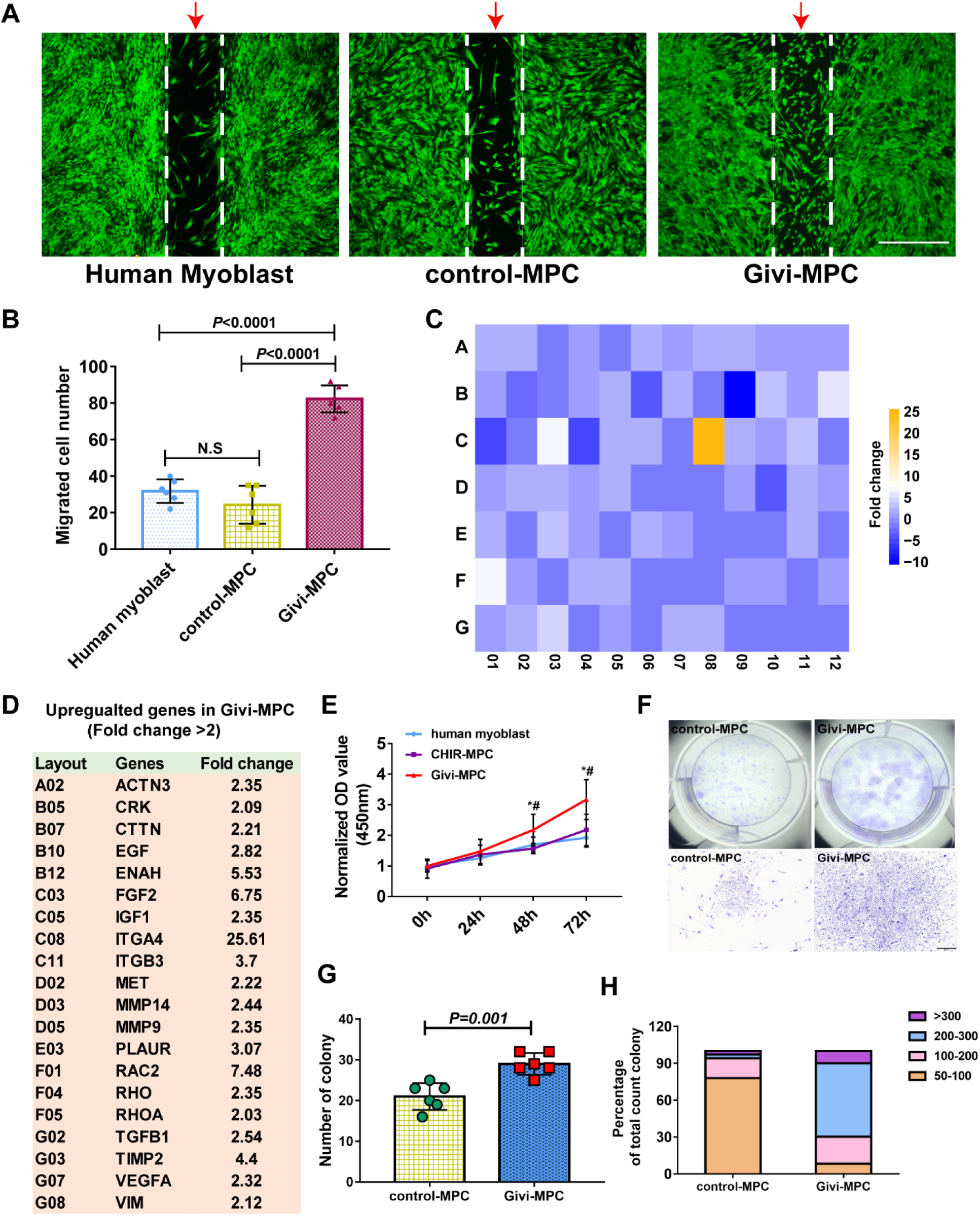
Givi-MPC exhibit superior proliferation and migration capacity. (A) Representative images and quantitative estimate (b) of cell migration by adult human myoblasts, and control-MPC, Givi-MPC (arrow). Cells were stained with Calcein AM (green). Bar = 1 mm. (B) Quantitative estimate of migrated cells. Givi MPC showed highest number of cells migrated compared with human myoblasts (*P<0.0001*) or CHIR99021 induced MPC (*P<0.0001*). No significant difference was observed between human myoblasts and control-MPC. (C) Heat map of the Human RT^2^ motility PCR Array. (D) Upregulated migration related genes in Givi-MPC vs. control-MPC using human cell motility PCR array. (E) The proliferation curves of human myoblasts vs MPC using CCK-8 assay. \**P< 0.05*; ^#^*P<0.05* vs control-MPC. n = 6. (F) Morphology of MPC colony. Bar = 500 μm. Number of colonies (G) and percentage of colonies with different cell number (H). control-MPC: CHIR99021 induced MPC; Givi-MPC: CHIR99021 and Givinostat induced MPC.

### *In vivo* engraftment of Givi-MPC restores dystrophin and integrated into the recipient environment

We transplanted human myoblast, control-MPC and Givi-MPC into Mdx/SCID mice with CTX injury, respectively. One-month post-transplant, Givi-MPC showed increased engraftment capacity and restoration of dystrophin than treatment with control-MPC and human myoblasts (**Fig.4A** **and 4B**). **Fig.S2** showed the engrafted Givi-MPC (GFP positive) expressed dystrophin. Quantitative data showed Givi-MPC treated TA muscle had significantly higher number of dystrophin positive muscle fibers (**Fig.4C**) and GFP and human laminin double positive muscle fibers (**Fig.4D**). To determine the functionality of the newly formed muscle fibers from Givi-MPC, we tested whether they were integrated into the recipient environment with innervation. Positive staining of presynaptic marker α-BTX was observed in close proximity to dystrophin positive muscle fibers in Givi-MPC treated TA muscle, suggesting the presence of neuromuscular junction in these muscle fibers (**Fig.4E**).

**Fig. 4.**
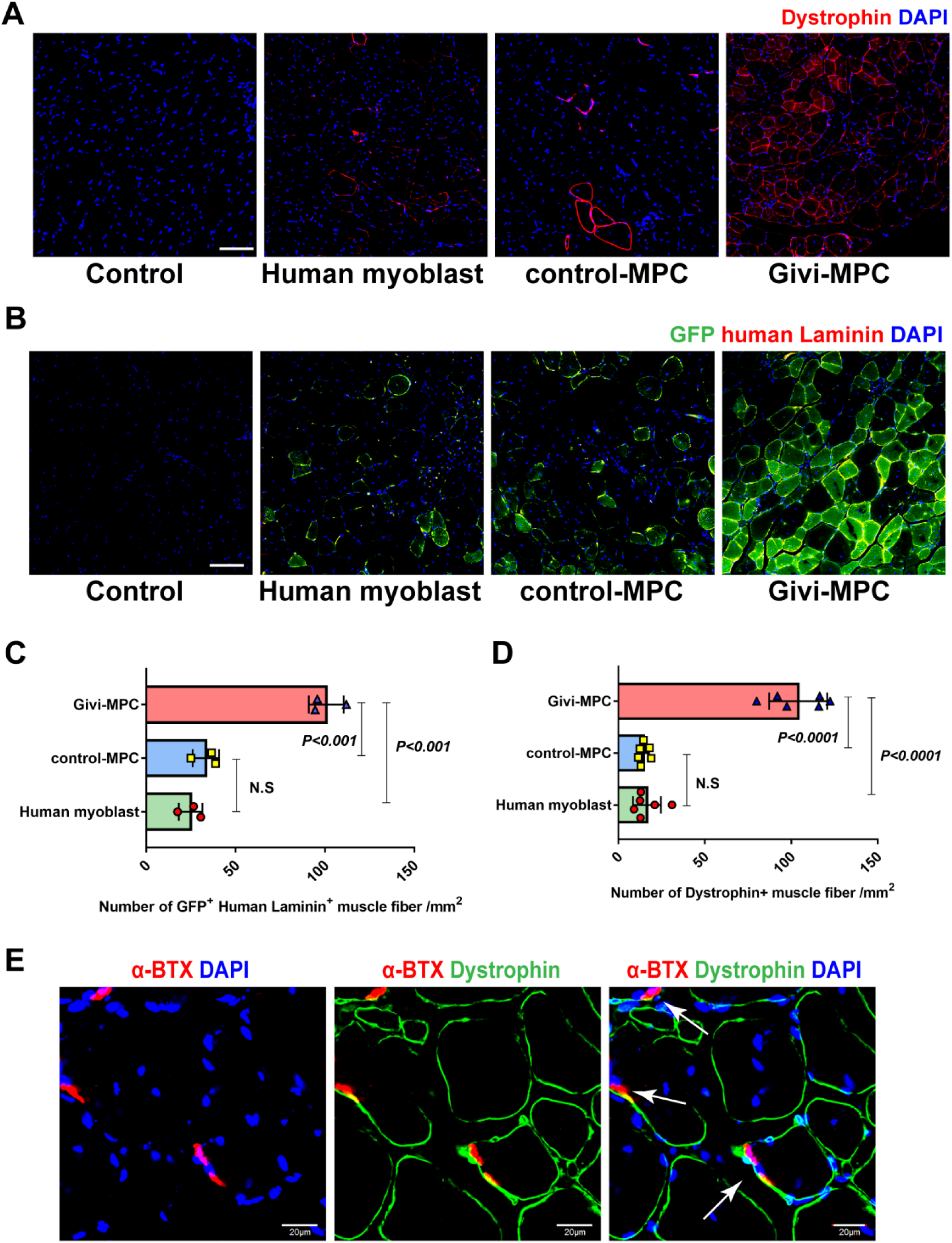
In *vivo* myogenic potential of different MPC and myoblast in Mdx/SCID mice with CTX injury. (A) Dystrophin restoration in Mdx/SCID mice by MPC transplantation at 1M after CTX injury. Bar = 50 µm. (B). Transplanted cells were labeled with GFP (Green) and identified with human laminin staining (Red). Quantitation of engrafted fibers at 1M: Dystrophin^+^ fibers (n = 6) (C) and human laminin and GFP double positive fibers (n = 3) (D). (E) Cross-section showing pre-synaptic staining with α-bungarotoxin in dystrophin positive fibers (n = 3). Bar = 20 µm.

### Givi-MPC limited inflammation, muscle necrosis and reduced fibrosis in Mdx/SCID mice post CTX injury

Hematoxylin and eosin and trichrome Masson staining revealed infiltration of inflammatory cells, and necrotic muscle fibers in Mdx/SCID mice 7 days’ post CTX injury (**Fig.5A**). A significant decrease in muscle necrosis was observed in Givi-MPC treated TA muscle compared to collateral PBS treated TA muscle (**Fig.5B**). Amongst different MPCs which were transplanted, Givi-MPC reduced muscle necrosis the most (**Fig.5C**) with reduced number of CD68 positive macrophages as compared with human myoblast and control-MPC treated tissue (**Fig.5D, E**) 7 days’ post CTX injury. In the muscle following 1M post CTX injury, transplantation of human myoblasts, control-MPC and Givi-MPC significantly decreased muscle necrosis compared to PBS treated collateral TA muscle (**Fig.6A-6D**). No significant difference in muscle fiber necrosis was observed between human myoblasts and control-MPC treated TA muscle. However, Givi-MPC treatment resulted in reduced necrosis area compared to other MPCs treatment (**Fig.6E**). Similarly, Givi-MPC transplantation reduced collagen deposits (red) compared to PBS, human myoblasts and control-MPC treated muscle (**Fig.6F, 6G-6I**).

**Fig. 5.**
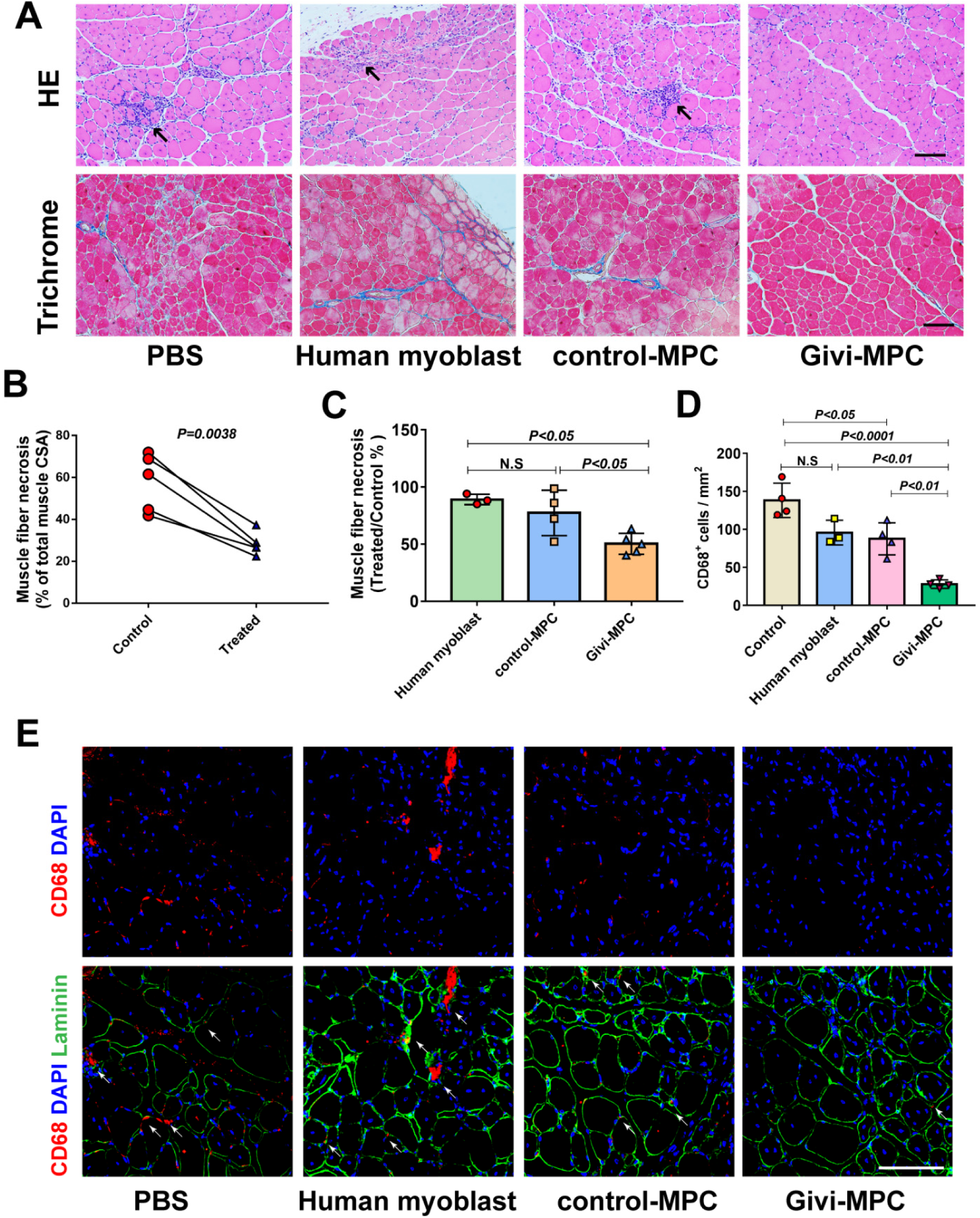
Givi-MPC decrease inflammation and muscle necrosis in Mdx/SCID mice 7 days after CTX injury. (A) Representative images of HE and Trichrome Masson staining in Mdx/SCID mice with human myoblasts or control-MPC or Givi-MPC transplantation 7 days after CTX injury. Black arrows indicate infiltrated inflammatory cells. (B) Quantification of muscle fiber necrosis between PBS treated collateral TA muscle or Givi-MPC treated TA muscle 7 days after CTX injury. (C) Quantification of muscle fiber necrosis of TA muscle among human myoblast or control-MPC or Givi-MPC transplantation mice 7 days after CTX injury. (D) Quantification of CD68 positive cells in TA muscle following MPC transplantation 7 days after CTX injury. (E) Representative images of macrophages (red, CD68) and human cells in TA muscle of Mdx/SCID mice with CTX injury following MPC transplantation.

**Fig. 6.**
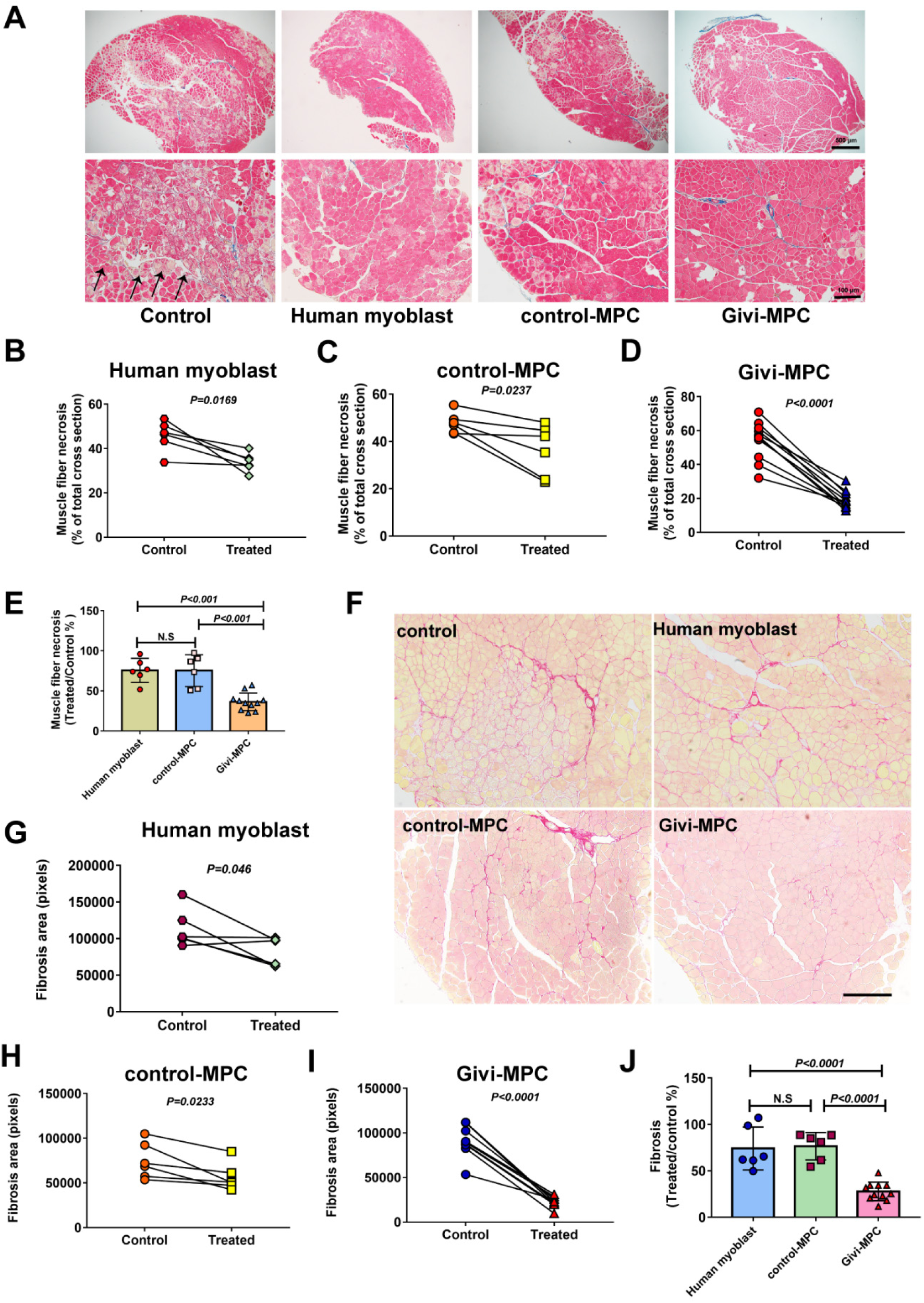
Givi-MPC decrease muscle necrosis and fibrosis in Mdx/SCID mice 1M after CTX injury. (A) Representative images of HE and Trichrome Masson staining in Mdx/SCID mice after transplantation with human myoblasts or control-MPC or Givi-MPC 1M after CTX injury. Bar = 500 μm (4×) and Bar = 100 μm (20×). Quantification of necrotic muscle fibers after treatment with human myoblasts (B), control-MPC (C) and Givi-MPC (D) 1M after CTX injury. (E) Comparison of muscle necrosis among human myoblasts or control-MPC or Givi-MPC transplantation mdx/SCID mice. (F) Representative images of tissue stained with Sirius red from Mdx/SCID mice. Bar = 100 µm. Quantification of muscle fiber fibrosis in collateral TA muscle treated with human myoblasts (G), control-MPC (H) and Givi-MPC (I) 1M after CTX injury. (J) Muscle fibrosis after transplantation of different MPCs in mdx/SCID mice.

### Givi-MPC repopulated the muscle stem cell pool

A significant number of Givi-MPC were transformed into muscle stem cells and occupied their sites as evidenced by double positivity for Pax7 and HNA cell under basal lamina at 1M post-transplantation (**Fig.7A**). A schematic outline of reinjury experiments with CTX is provided (**Fig.7B**). Compared with contralateral PBS treated TA muscle, expression of dystrophin was detected in Givi-MPC treated TA muscle after reinjury (**Fig.7C**). Furthermore, Givi-MPC treated TA muscle showed increased muscle regeneration and less infiltration of inflammatory cells compared with contralateral PBS treated muscle (**Fig.7D**). These data indicated that the engrafted Pax7 positive cells responded to reinjury and formed new muscle fibers.

**Fig. 7.**
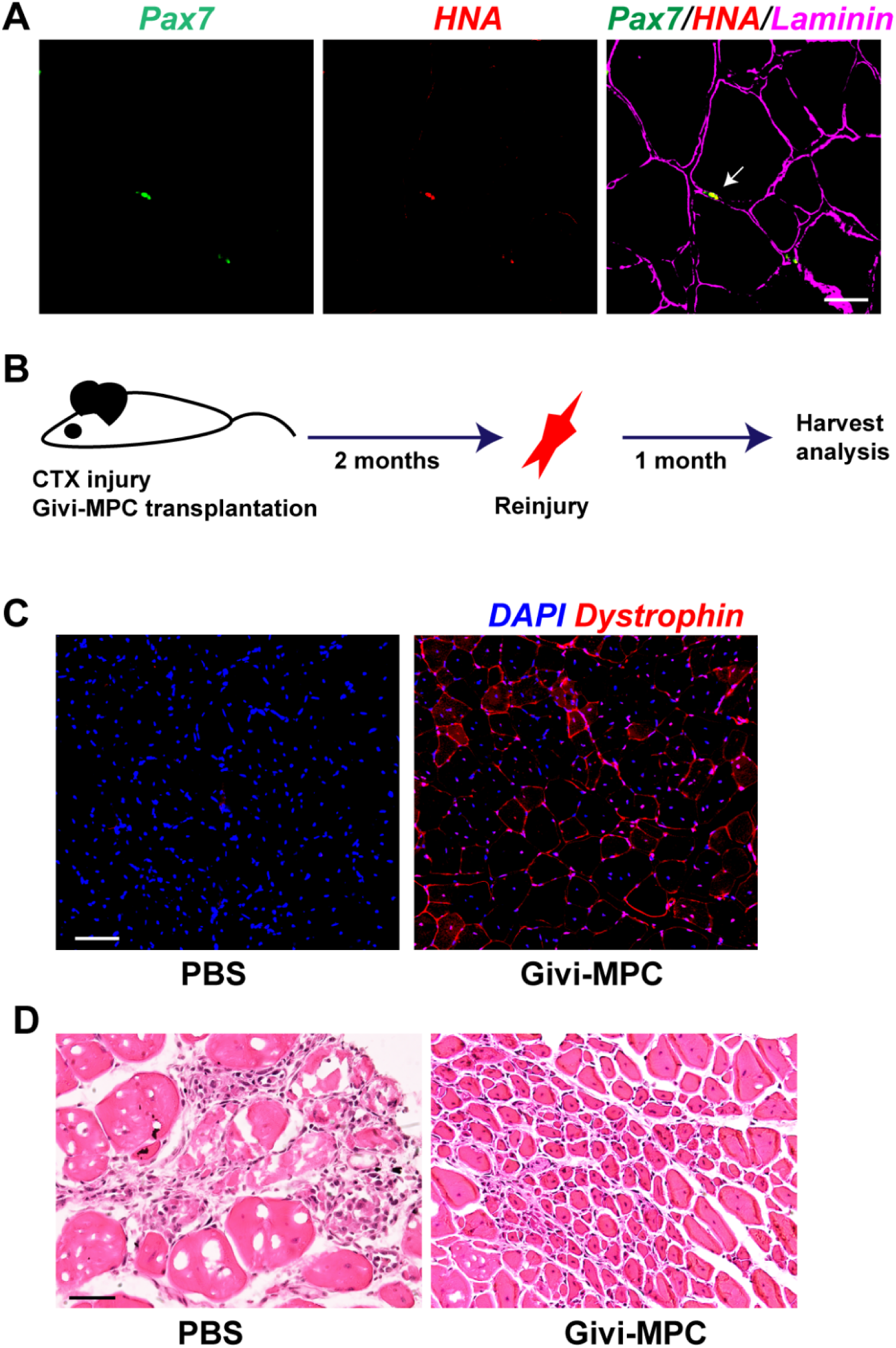
Givi-MPC repopulated the muscle stem cell pool. (A) Muscle cells positive for Pax7 (green) and human nuclear antigen (red) cell under the basal lamina from Mdx/SCID mice after 1M of Givi-MPC transplantation. Bar = 20 µm. (B) Schematic of reinjury experiment. (C) 1M after reinjury, expression of dystrophin in Givi-MPC treated TA muscle tissue and contralateral PBS treated TA muscle tissue. Bar = 50 µm. (D) Representative HE stained images of Givi-MPC treated TA muscle tissue and contralateral PBS treated TA muscle tissue. Bar = 50 µm.

### Extracellular vesicles derived from Givi-MPC facilitated angiogenesis in muscle following CTX injury

Angiogenesis is critical for muscle regeneration (18, 19). Givi-MPC treatment caused higher capillary density (CD31 positivity) in TA muscle 1M post CTX injury (**Fig.8A** **and 8B**). Next we tested whether increased angiogenesis was due to paracrine effects by EV released from MPCs. We isolated EV from Givi-MPC using size exclusion columns. Using tunable resistive pulse sensing (TRPS) technique, we measured the concentration and size of EV from Givi-MPC. The size of isolated EV was roughly 118 ± 31.7 nm (**Fig.S3A, B**). *In vitro* tube formation assay indicated that EV from Givi-MPC promoted tube formation (**Fig.8C**) and resulted in higher average tube length (**Fig.8D**) compared to treatment with EV from human myoblasts or control-MPC. We further analyzed the miRNA cargo contents of EV from Givi-MPC. A heatmap of significantly upregulated and downregulated miRNAs in EV from Givi-MPC compared to EV from human myoblasts was shown in **Fig.8E**. miR-210, miR-181a, miR-17 and miR-107 were enriched in EV from Givi-MPC.

**Fig. 8.**
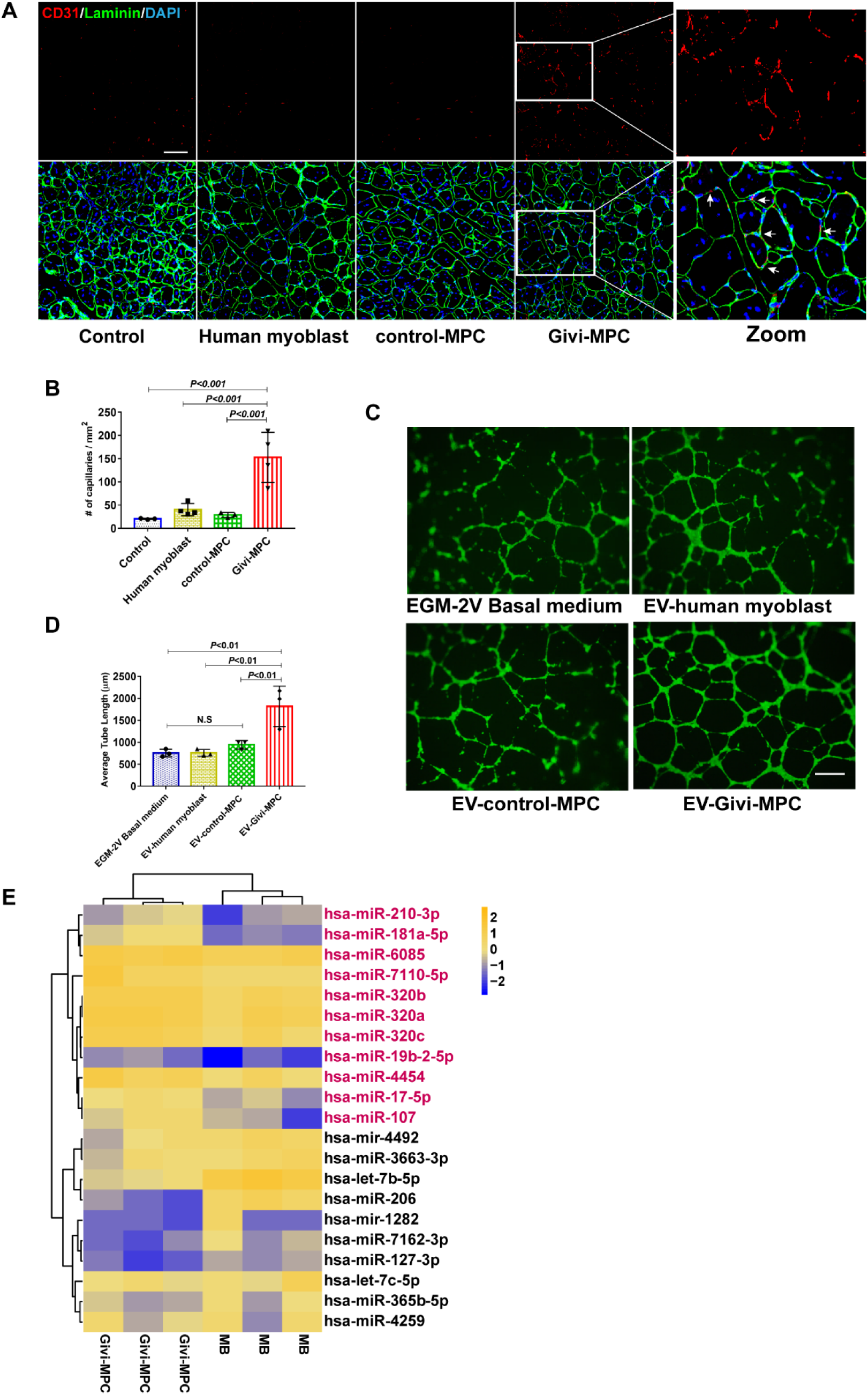
Extracellular vesicles derived from Givi-MPC promoted angiogenesis. (A) Representative images of CD31 (Red) and laminin (Green) staining in Mdx/SCID mice 1M post injury. Bar = 50 µm. (B) Quantification of capillary density (CD31 positive capillaries). (C) Representative images of tube formation by human aortic endothelia cells (HAECs) following EV treatment from human myoblasts, or control-MPC or Givi-MPC (1μg/well, 24 well plate). HAECs were labeled with Calcein AM (Green). Bar = 500 μm. (D) Tube formation assay. Average tube length was analyzed from 3 biological repeated experiments. (E) Heatmap showing significant upregulation of miRs in EV derived from Givi-MPC compared to EV-human myoblasts.

## Discussion

In the present study, we successfully generated highly proliferative and integration free MPC from multiple hiPS cell lines using CHIR99021 and Givi. These cells expressed myogenic markers including Pax7 and desmin, which were also capable to differentiate into muscle cells under specific differentiation medium in *vitro*. Of particular significance was the ability of these MPCs to differentiate in dystrophic mouse model, making them more suitable for therapeutic applications. These cells possess special properties which make them unique for therapeutic applications. Migration and engraftment of transplanted cells to the site of injury are crucial to initiate differentiation into skeletal muscle components in the dystrophic muscle(20, 21). Limited cell migration hampers engraftment efficiency in skeletal muscle (22, 23). In the present study, we found MPC induced by Givi exhibited superior migration and proliferation capabilities compared with human myoblasts and control MPC generated by CHIR99021 and FGF. Go analysis further showed upregulation of cell migration related genes enabling them to migrate to distant injured muscle (10). In our data, genes related to migration were significantly upregulated with Givi treatment. ITGA4 was the most upregulated gene with 25.61-fold change. Integrin subunit α4 (ITGA4) is a member of the integrin alpha chain family of proteins. Integrin α subunits which pair with β1 play a critical role during *in vivo* myogenesis. Integrin α4 subunit is expressed in the myotome and in early limb muscle masses during muscle development (24, 25). Murine Lbax1^+^ embryonic muscle progenitors expressed ITGA4 (26). It has been reported that teratoma derived MPC possessed high engraftment efficiency in muscle dystrophy model (10). However, the mechanism of upregulation of ITGA4 by Givi and migration medicated by ITGA4 need further study. DMD is a disease with body-wide systemic and progressive skeletal muscle loss, thus further study for the role and mechanism of ITGA4 in MPC migration will move MPC-based therapy for DMD forward to clinical application. In agreement with *in vitro* observations, we also observed higher engraftment efficiency of Givi-MPC compared to human myoblasts and control MPC upon transplantation in muscle tissue from Mdx/SCID mice following CTX injury. The significant engraftment in muscles of Mdx/SCID mice by human iPS-derived skeletal myogenic progenitors resulted in more dystrophin expressing myofibers or human laminin positive myofibers. Besides dystrophin, presence of neuromuscular junctions in human myofibers using α-BTX together with dystrophin in Mdx/SCID mice with Givi-MPC transplantation, suggest that formation of functional myofibers has occurred.

Histological analysis showed that fewer muscle fibers had undergone necrosis and fibrosis in injured TA muscle of Mdx/SCID mice treated with Givi-MPC. Inflammatory cell infiltration in general contributes to myofiber necrosis (27, 28). Although Mdx/SCID mice are immunodeficient, it has been reported that M1 macrophages participated in skeletal muscle regeneration in SCID mice (29), suggesting partial immune reactivity in these mice. It has been reported Givi has potential anti-inflammatory effects (30, 31). For example, Givi decreased inflammation in a mouse model with myocardial infarction (31). With HE staining, we found infiltration of larger number of inflammatory cells in TA muscle from Mdx/SCID mice treated with PBS, or human myoblasts or control MPC treatments 7 days post-CTX injury. Negligible macrophage infiltration identified by CD68 staining was observed in Givi-MPC transplanted Mdx/SCID mice 7 day post-CTX injury. These observations support that Givi-MPC had anti-inflammatory effects upon transplantation in CTX injured muscle suggesting that properties of MPC depend on the source of reprogramming molecule. Besides immediate effects on engraftment and differentiation, the long-term maintenance of newly formed skeletal muscle is ultimately dependent on the ability of the transplanted MPCs to contribute to the skeletal muscle stem cell pool (10). Here we observed Givi-MPC derived Pax7 positive cells under basal lamina upon transplantation, and with subsequent reinjury the Givi-MPC contributed to secondary regeneration in the Mdx/SCID mice. This observation supported that a subpopulation of Givi-MPC can seed the stem cell pool.

Angiogenic impairment of the vascular endothelial cells (EC) isolated from mdx mice compared with wild type mice has been reported (32) causing a marked decrease in the vasculature in TA muscle of mdx mice (33). Local delivery of muscle-derived stem cells engineered to overexpress human VEGF into the gastrocnemius muscle of Mdx/SCID mice resulted in marked increase in angiogenesis accompanied by enhanced muscle regeneration and decreased fibrosis compared with mice treated with non-engineered cells (34). In addition, satellite cells isolated from mdx mice exhibited reduced capacity to promote angiogenesis, as demonstrated in a co-culture model of satellite cells of Mdx mice and microvascular fragments (35). Here, our study demonstrated that after Givi-MPC transplantation, an increase in capillary density was observed as evidenced by CD31 staining in CTX injured Mdx/SCID mice compared to treatment with other MPCs. These results enforce the idea that an interaction between EC and MPC was important for myogenesis and angiogenesis *in vitro* and *in vivo* during skeletal muscle regeneration (18). To further strengthen this observation, we found that EV from Givi-iMPC were enriched in several miRNAs including miR-181a, miR-17, miR-210 and miR-107, miR-19b compared with EV from human myoblasts. Due to role of EV in cell-to-cell communication, these enriched miRNAs have been demonstrated to participate in angiogenesis. Activation of miR-17-92 cluster promoted angiogenesis via PTEN signaling pathway, however, EC miR-17-92 cluster knockout impaired angiogenesis (36). miR-181a and miR-210 are also reported to promote angiogenesis (37–40). Thus it is very likely that Givi-MPC interacted with resident EC to initiate myogenesis and angiogenesis in Mdx/SCID mice after CTX injury.

## Conclusion

We successfully generated highly expandable and integration free MPC from multiple hiPS cell lines using CHIR99021 and givinostat. Givinostat induced MPC were highly proliferative and migratory and transplantation resulted in marked and impressive myoangiogenesis and restored dystrophin in injured TA muscle compared to control MPC and adult human myoblasts. It is concluded that hiPSCs pharmacologically reprogrammed into MPC with a small molecule, givinostat with anti-oxidative, anti-inflammatory and muscle gene promoting properties is an effective cellular source for treatment of muscle injury and restoration of dystrophin in DMD.

