## Supplemental figures for "Human iPS Cells Derived Skeletal Muscle Progenitor Cells Promote Myoangiogenesis and Restore Dystrophin in Duchenne Muscular Dystrophic Mice"

Fig.S1 The schematic outline for control-MPC generation.

Fig.S2 Engrafted Givi-MPC (GFP positive) expressed dystrophin. Bar = 100 µm.

Fig.S3 (A) Extracellular vesicles (EV) isolated from Givi-MPC visualized by transmission electron microscopy (TEM). (B) The size of isolated EV from Givi-MPC was roughly 118 ± 31.7 nm.

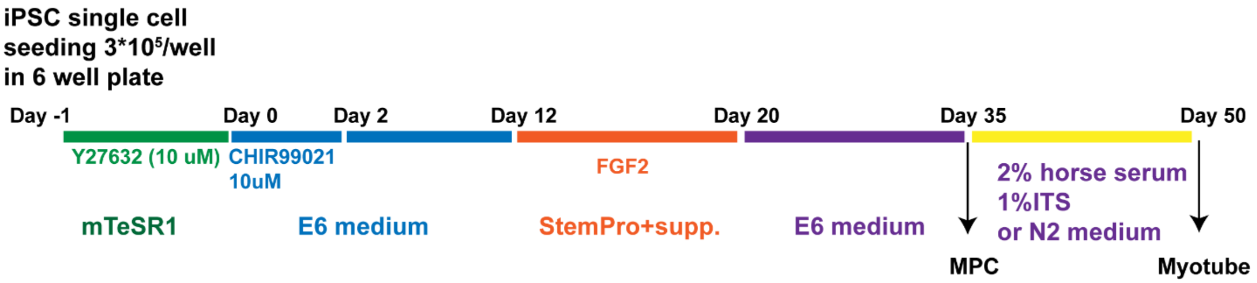


Fig.S1

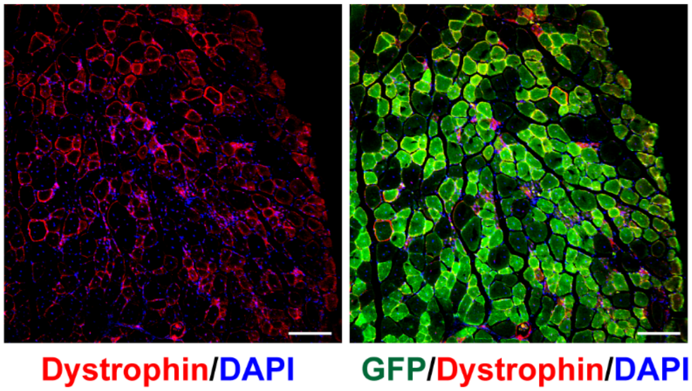


Fig.S2

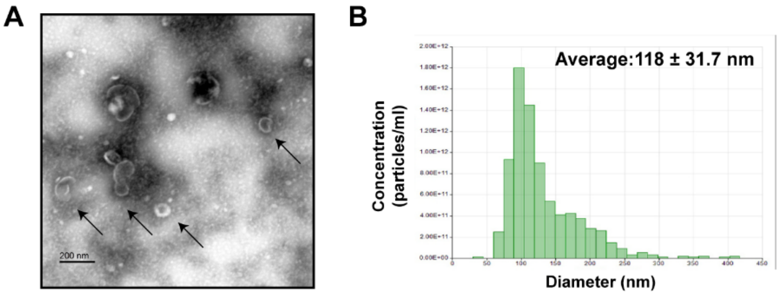


Fig.S3
